# The influence of heteroresistance on minimum inhibitory concentration (MIC), investigated using weak-acid stress in food spoilage yeasts

**DOI:** 10.1101/2023.02.06.527412

**Authors:** Joseph Violet, Joost Smid, Annemarie Pielaat, Jan-Willem Sanders, Simon V. Avery

## Abstract

Populations of microbial cells may resist environmental stress by maintaining a high population-median resistance (IC_50_) or, potentially, a high variability in resistance between individual cells (heteroresistance); where heteroresistance would allow certain cells to resist high stress, provided the population was sufficiently large to include resistant cells. This study sets out to test the hypothesis that both IC_50_ and heteroresistance may contribute to conventional minimal-inhibitory-concentration (MIC) determinations, using the example of spoilage-yeast resistance to the preservative sorbic acid. Across a panel of 26 diverse yeast species, both heteroresistance and particularly IC_50_ were positively correlated with predicted MIC. A focused panel of 29 different isolates of a particular spoilage yeast was also examined (isolates previously recorded as *Zygosaccharomyces bailii*, but genome resequencing revealing that several were in fact hybrid species, *Z. parabailii and Z. pseudobailii*). Applying a novel high-throughput assay for heteroresistance, it was found that IC_50_ but not heteroresistance was positively correlated with predicted MIC when considered across all isolates of this panel, but the heteroresistance-MIC interaction differed for the individual *Zygosaccharomyces* subspecies. *Z. pseudobailii* exhibited higher heteroresistance than *Z. parabailii* whereas the reverse was true for IC_50_, suggesting possible alternative strategies for achieving high MIC between subspecies. This work highlights the limitations of conventional MIC measurements due to the effect of heteroresistance in certain organisms, as the measured resistance can vary markedly with population (inoculum) size.

**Importance:** Food spoilage by fungi is a leading cause of food waste, with specialised food spoilage yeasts capable of growth at preservative concentrations above the legal limit, in part due to heteroresistance allowing small subpopulations of cells to exhibit extreme preservative resistance. Whereas heteroresistance has been characterised in numerous ecological contexts, measuring this phenotype systematically and assessing its importance are not encompassed by conventional assay methods. The development here of a high-throughput method for measuring heteroresistance, amenable to automation, addresses this issue and has enabled characterisation of the contribution that heteroresistance may make to conventional MIC measurements. We used the example of sorbic acid heteroresistance in spoilage yeasts like *Zygosaccharomyces* spp, but the approach is relevant to other fungi and other inhibitors, including antifungals. The work shows how median resistance, heteroresistance and inoculum size should all be considered when selecting appropriate inhibitor doses in real-world antimicrobial applications such as food preservation.

## Introduction

Resistance of microorganisms to inhibitory agents is a growing problem in therapeutics (Andersson et al., 2019, Dewachter et al., 2019) and food supply chains (Davies et al. 2020). The resistance of a microorganism to a stressor is typically described using a conventional minimum inhibitory concentration (MIC) measurement, which reflects the concentration of a stressor required to prevent growth at a particular fixed inoculum size (Box 1). Other measures of stress resistance include IC_50_, a population-median measure describing the concentration required to inhibit either 50% of cells in a population (as used in the present work, Box 1) or all cells by 50%, depending on the study. In recent years there has been growing recognition also of heteroresistance, describing the heterogeneity of resistance between individual clonal cells (Box 1), which appears to be very ubiquitous (Bishop et al. 2007, Holland et al. 2014, Jing et al. 2018, Levy et al., 2012). However, the extent to which differences in heteroresistance may contribute to MIC of a cell inoculum remains barely explored, especially relative to traditional measures such as IC_50_. The present work aimed to address this gap in our knowledge, as it could explain variability in MIC data that are used as common standards, due to parameters such as population size and the inoculum effect (Box 1).

### Box 1: Definitions of parameters

1. **MIC:** minimum inhibitory concentration, i.e., lowest concentration of a stressor required to completely inhibit the growth of an inoculum of a microorganism.
2. **MIC^*EXP*^:** Experimentally derived MIC, at a given inoculum size, e.g., the MIC^*EXP*^ for 100 cells of a given strain could be different to the MIC^*EXP*^ for 10,000 cells of the same strain.
3. **MIC^*MODEL*^:** MIC of 10,000 cells/well predicted using a lognormal distribution curve, fitted to MIC^*EXP*^ values (see *Fitting to normal and lognormal distributions*)
4. **Heteroresistance:** Cell-cell phenotypic heterogeneity in a resistance phenotype. Here quantified as the standard deviation of a lognormal distribution of single cell resistances
5. **IC_50_:** Population-median resistance, here defined as the mean of a lognormal distribution of single cell resistances
6. **Inoculum effect:** Phenomenon by which MIC^*EXP*^ increases with inoculum size, due to increased probability of resistant cells being present

Phenotypic heterogeneity among individual cells of genetically uniform populations is considered an evolutionarily selected strategy akin to bet hedging (Ackermann 2015). It describes cell-to-cell variation in a phenotype (e.g. stress resistance) within a cell population, resulting in phenotypically (not genotypically-) distinct subpopulations among which some may be better equipped to thrive if conditions change (Ackermann 2015, Smith et al., 2007, Levy et al. 2012, Stratford et al 2019). The rationale is that generation of phenotypic heterogeneity, may be a relatively low cost strategy (metabolically) for improving survival-chances in uncertain conditions. Such heterogeneity in the case of stress resistance can be observed in almost any resistance phenotype: examples of heteroresistance are evident in microbial responses to antibiotics (Dewachter et al., 2019, Scheler et al., 2020, Jing et al. 2018), heat (Levy et al., 2012), osmotic (Stratford et al. 2019), metal (Holland et al., 2014) and weak organic acid (Stratford et al. 2014, Geoghegan et al. 2020) stresses. Mechanistic determinants of heteroresistance have been characterised, including cell-to-cell variation in gene expression linked to particular ‘high’ or ‘low’-variation promoters or transcription factors, commonly referred to as gene expression noise (Silander et al. 2011, Urchueguia et al., 2021, Newman et al. 2006). Housekeeping genes tend to be expressed with low variation between cells, whereas high-variation promoters are more commonly associated with stress-response genes (Newman et al. 2006). Engineering increased variation of expression of an oxidative stress response gene has been shown to increase resistance to oxidative stress in *Saccharomyces cerevisiae* (Liu et al., 2018), while dampening of expression variation in similar functions gave decreased resistance (Smith et al., 2007).

As food spoilage microorganisms can exhibit both high population-median resistance and heteroresistance to food preservatives (Stratford et al. 2014), they provide a good example with which to address this study’s main aim to characterise the relationship between MIC and heteroresistance versus population-median resistance. Among the major food preservatives, sorbic acid is a weak organic acid preservative widely used in condiments and drinks of pH ≤4, conditions in which the acid is undissociated and capable of traversing the plasma membrane. Several modes of action have been proposed for sorbic acid, including cytosolic acidification (Stratford et al., 2013a) and disruption of the mitochondrial electron transport chain (Stratford et al., 2020). Efficacy of sorbic acid as a food preservative is hampered by the intrinsic resistance of specialised spoilage yeasts, which are capable of growth at sorbic concentrations exceeding the maximum legal limit dictated by food standards agencies (Stratford et al., 2013b).

*Zygosaccharomyces bailii* is a food spoilage yeast exhibiting a high level of resistance to weak organic acids (Stratford et al. 2014). Two interspecies hybrids related to *Z. bailii* and also reported to spoil preserved foods are *Z. parabailii* and *Z. pseudobailii*, formed through hybridisation events between ancestral *Z. bailii* and two unknown *Zygosaccharomyces* species, sharing approximately 90-93% sequence identity with *Z. bailii* (Braun-Galleani et al., 2018). *Z. bailii* has several proposed mechanisms of resistance to weak organic acids, including a relatively thick and rigid plasma membrane (Lindberg et al. 2013, Lindahl et al. 2016), metabolism of sorbic acid to less inhibitory products (Mollapour and Piper, 2001), metabolism that is primarily fermentative (Stratford et al. 2020) and a low cytosolic pH (Stratford et al. 2013b, 2014). In addition to being relatively resistant to sorbic acid (according to MIC or IC_50_), *Z. bailii* also shows high sorbic acid heteroresistance (Stratford et al. 2013b), with a small proportion of cells capable of resisting extreme concentrations even though they may take several weeks to grow to detectable levels.

The food-industry relevance of *Z. bailii*, coupled with its high heteroresistance makes it an ideal model for studying the relative contributions of population-median resistance and heteroresistance to MIC. In this study we assess and compare these parameters in a panel of diverse spoilage yeasts as well as more select panels of *Z. bailii, Z. parabailii* and *Z. pseudobailii* isolates. The work highlights the importance of context when assessing the resistance of a microorganism, with population-median resistance, heteroresistance and inoculum size impacting the resistance of a given inoculum, nuances which are lost when an MIC at a single inoculum size is taken.

## Materials and Methods

### Yeast species and strains

Yeast species and strains used for experiments in this study are listed in Tables 1, 2 (alternative identifiers and origin of isolation are given where available). Members of a large, initial *Zygosaccharomyces* panel (Table S1) were designated *Z. bailii, Z. parabailii* or *Z. parabailii* based on whole genome sequencing (WGS) analysis (see Genome sequencing) and a subset of these comprised the main *Zygosaccharomyces* panel used for experiments (Table 1). Several species previously designated *Z. bailii* were redesignated *Z. parabailii* or *Z. pseudobailii* based on the WGS analysis (below). Members of a wide panel of isolates from more diverse yeast genera (Table S2) were identified previously based on sequencing of the D1/D2 region of the 25S ribosomal DNA (rDNA) (Stratford et al. 2013b).

**Table 1:**
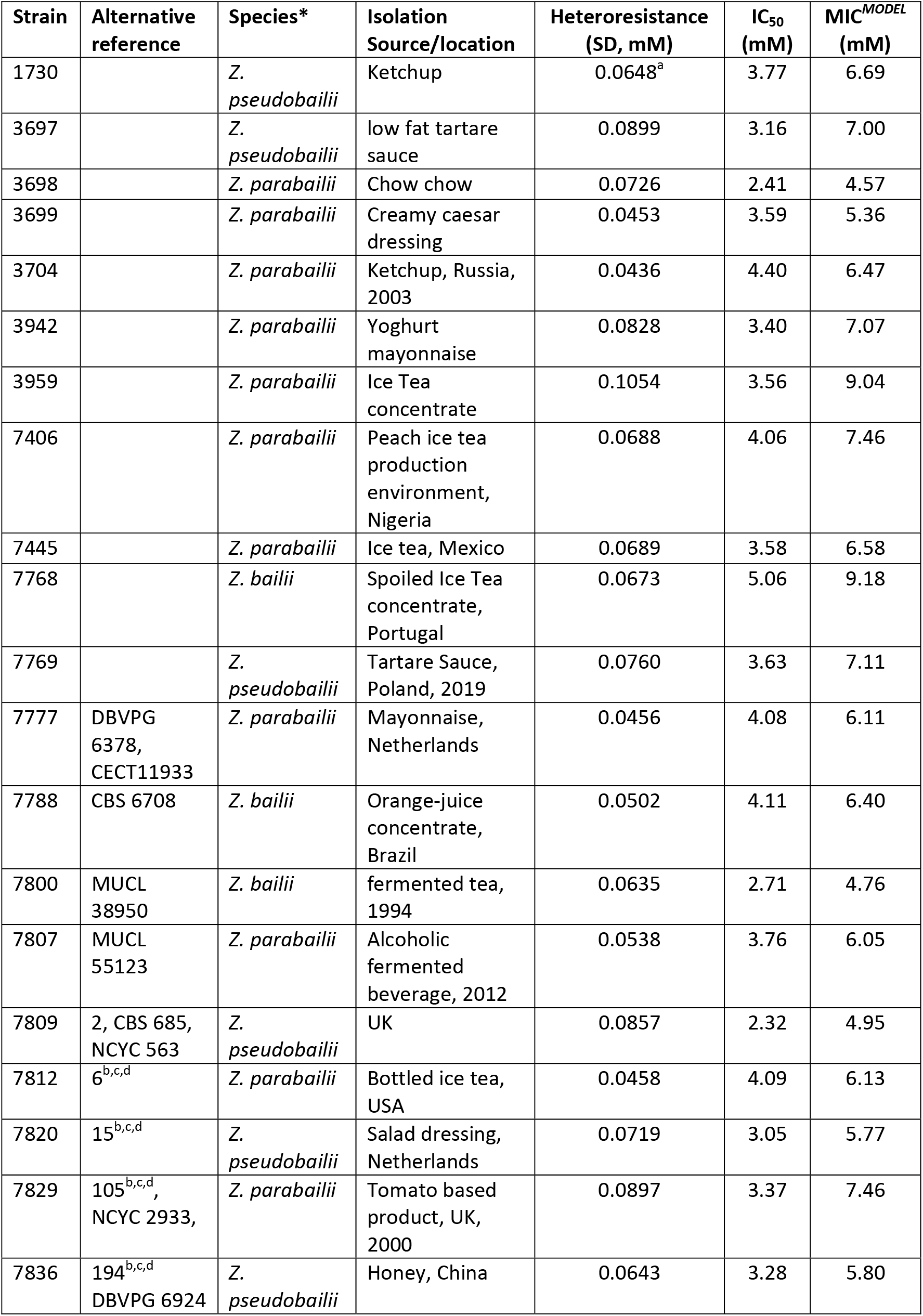
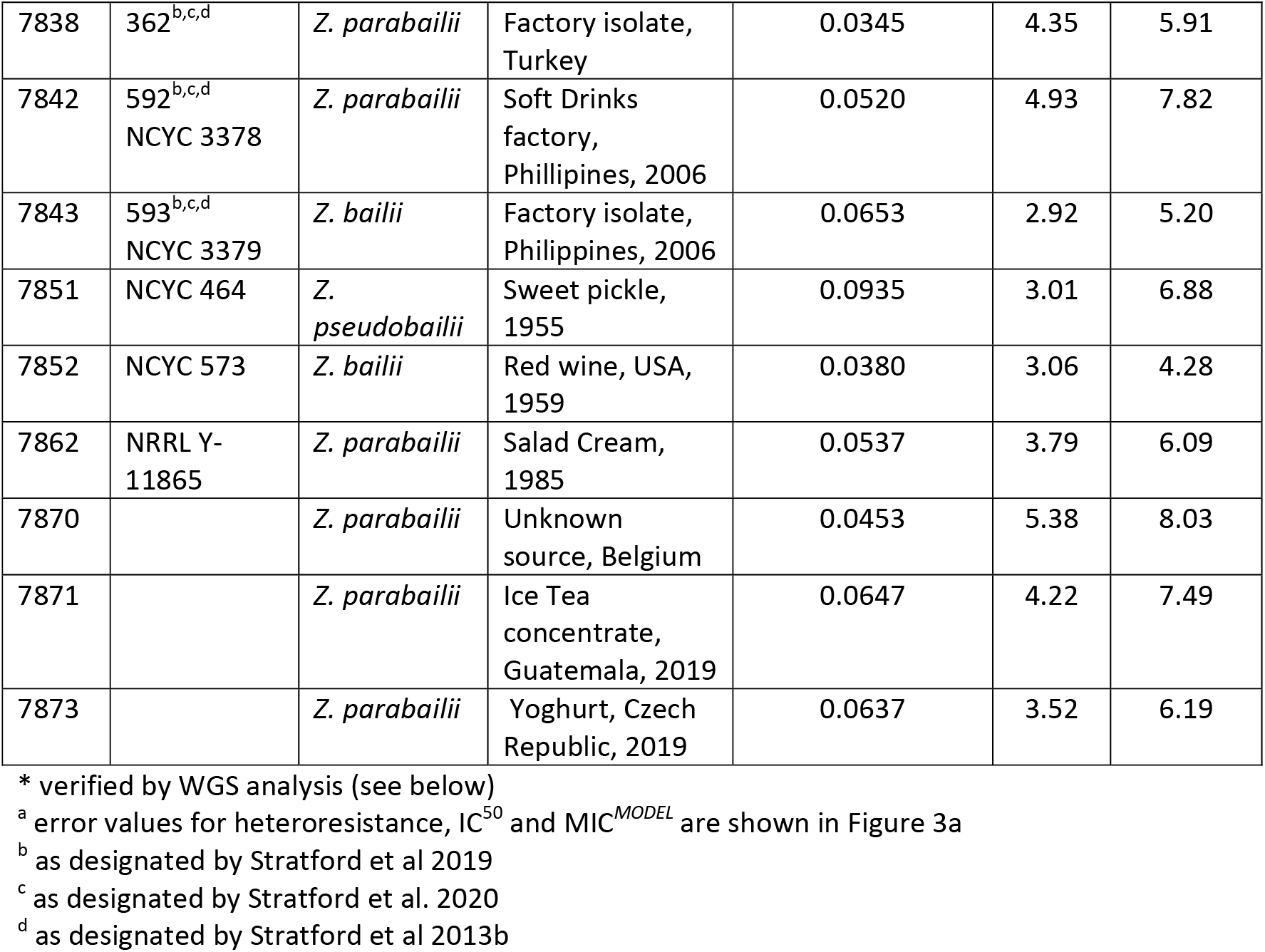
Main panel of *Zygosaccharomyces* strains, including sorbic acid resistance values.

### Culture conditions

The YEPD growth medium used for culturing [20 g/l glucose, 20 g/l bacteriological peptone (Oxoid), and 10 g/l yeast extract (Oxoid)] was adjusted to pH 4 using 5 M HCl prior to sterilisation by autoclaving. Yeasts were stored at −80°C in cryovials in YEPD medium mixed with 0.2 volumes glycerol. Cells were grown from frozen on YEPD supplemented with 2% agar (Oxoid) for 3 days at 30°C, before maintenance at 4°C for up to 3 weeks. For each biological replicate in experiments, a single colony was suspended in 1 ml YEPD broth and, after determination of OD_600_, an aliquot transferred to 10 ml YEPD broth in order to give a starting OD_600_~0.02. These cultures were incubated in 50 ml Erlenmeyer flasks at 24°C for >12 hours with shaking (120 rev/min) prior to use in experiments at OD_600_ 0.1 – 2.0, when cells were growing in exponential phase. Desired concentrations of sorbic acid in the growth medium were produced using a stock solution of 20 mM potassium sorbate (Sigma-Aldrich) in YEPD, adjusted to pH 4 using 5 M HCl. Final concentrations of sorbic acid in YEPD agar or broth were produced by mixing appropriate ratios of YEPD medium (pH 4) with the 20 mM sorbic acid-supplemented medium. All steps were carried out under aseptic conditions.

### *De novo* genome sequencing

*De novo* genome sequencing was carried out by KeyGene (Wageningen, Netherlands) by mapping short Illumina reads to long PacBio reads for *Z. bailii* (7846), *Z. parabailii* (3699, containing *Z. bailii*-derived ‘A’ genome and hybrid ‘B’ genome) and *Z. pseudobailii* (3697, containing *Z. bailii*-derived ‘A’ genome and hybrid ‘C’ genome). Yeast isolates were grown in YEPD medium at 25°C for 48 hours. Subsequently, cells were harvested by centrifugation (3000 *g*, 3 min), the supernatant removed and the pelleted cultures freeze dried prior to DNA isolation. Isolation of high molecular weight DNA suitable for long-read sequencing was carried out by Keygene based on methods described previously (Zhang et al. 2012, Datema et al. 2016).

To generate the sequencing data for *Z. pseudobailii* and *Z. parabailii*, one PacBio SMRT cell (Sequel) was used for each isolate. The same DNA was used for generating paired-end Illumina reads (MiSeq) that was used for polishing of the PacBio data. To obtain the initial assembly of the two hybrid species, *Z. parabailii* and *Z. pseudobailii*, the falcon-based HGAP 4 assembly method in SMRT Link portal v7.0 from Pacific Biosciences was used. As the method already includes the error correction of long reads, polishing was continued with Illumina short reads which has lower base calling error. The assembly was polished three times using Pilon software version 1.22. The inverted full size contigs in the assembly with respect to the reference genome were reverse complemented to make the assembly compatible with public reference. The contigs which were partially inverted were left as they were.

For the assembly of the *Z. bailii* genome, the sequencing library was prepared from PacBio high fidelity (HiFi) reads, with lower error rate (similar to Illumina short reads), hence no polishing was required. HiFi reads were obtained via circular consensus sequencing (CSS) from the SMRT Link portal. The reads were then assembled into the draft assembly using the HiFi option of the Canu assembler. The draft assembly was polished twice with HiFi reads using Racon software. Then, the repetitive haplotypes were cleaned using the Purge Haplotigs pipeline. The remaining contigs were checked for contamination by aligning the assembly to public reference, and by aligning a part of the contigs to the BLASTn database. Finally, for all final assemblies of the three *Zygosaccharomyces* subspecies, a BUSCO analysis was performed to check the completeness of genomes. The pairwise alignments of genomes were done via the nucmer algorithm from the Mummer 4 package and KeyGene’s proprietary STL aligner (multi genome aligner). The multiple genome alignments (as pan genomes) were visualized using Mauve software. All sequences used in this study were submitted to the ENA under the study code PRJEB59101; *de novo* sequences were assigned the accession numbers GCA_949129065, GCA_949129075 and GCA_949129085 for *Z. parabailii* (3699), *Z. bailii* (7846) and *Z. pseudobailii* (3697), respectively.

### Genome sequencing of the initial panel of *Zygosaccharomyces* isolates

The 111 isolates from the initial panel (Table S1) were cultured for 24 hours at 25°C in YEPD and genomic DNA (gDNA) was isolated using the CTAB method (Stirling et al. 2003). Some gDNA samples contained evidence of bacterial DNA, therefore, not all strains were included in the library preparation and resequencing. Resequencing was performed by Illumina HiSeq4000 with 2x 125 bp, on two lanes. For all Illumina sequencing, quality control was based on the QC30 threshold (80% of reads with ≤0.1% chance of miscalled base in all lanes of each run). The raw sequence data were trimmed on a minimum base quality of 30 (Phred scores) and sequences with one or more N-nucleotides were discarded. After trimming and filtering, read pairs for which both pairs had a minimum read length of 50 nt were kept for further analysis. The pre-processed sequencing data was mapped to the reference genome using BWA mem 0.7.17. Duplicated reads in the BAM file were marked by Picard-tools MarkDuplicates (v1.63). SAMTOOLS (v1.3) was used to view, index, sort and to merge the alignment files and the BAM files per sample genotyped using GATK 4. Variants such as SNPs and INDELs were identified using GATK4 Haplotypecaller and were stored in a single Variant Call Format (*.vcf) file. These variants were filtered on allele quality, sample quality and allele depth. A minimum allele quality of 30, minimum sample quality of 20 and minimum allele coverage depth of 7X were used for filtering. SNPs found in all samples that are identical were discarded as they reflect errors in the reference sequence. The filtered variants were annotated using the gene models with SNPeff. Most genes in *Z. parabailii* (3699) and *Z. pseudobailii* (3697) were assigned to the A-subgenome (highly similar to the *Z. bailii* 7846 genome) or to the other subgenome (i.e. B or C; see below). The long sequence aligner Blasr v1.3 was used in the Bidirectional best hit approach to identify homologs. Only coding sequences were used for this approach, extracted from the annotated gene models. Homologous genes with a high sequence similarity (i.e. above 98%) to *Z. baillii* genes were assigned to the A-subgenome and genes below the sequence similarity threshold were assigned to the B-subgenome (hybrid genome of *Z. parabailii*) or C subgenome (hybrid genome of *Z. pseudobailii*). Phylogenetic trees were constructed based on Genetic Distance Analysis of all open reading frames of sequences assigned to the A genome. The A genome of each sequence was compared to the A genome of *Z. parabailii* reference strain 3699 and an Unweighted pair group method with the given arithmetic mean (UPGMA) score (Khan et al., 2008) assigned, from which a phylogenetic tree was constructed (Figure S5). All sequences used in this study were submitted to the ENA under the study code PRJEB59101; strain-specific sample numbers are quoted in Table S1.

### Dose-response curve assay for heteroresistance

Yeasts were grown to exponential phase as described above and diluted in YEPD broth to OD_600_ ~0.02 (inoculum size in 75 μl ~10,000 cells) or OD_600_ 0.0002 (inoculum size in 75 μl ~100 cells). The use of the different inocula allowed reliable determination of % viability across a wider range of sorbic acid concentrations. Aliquots (75 μl) of each inoculum size were spread across 90 mm diameter Petri dishes containing 20 ml YEPD agar supplemented with different concentrations of sorbic acid (see above). After 21 days static incubation at 23°C, colony forming units (CFU) were enumerated and percentage survival calculated by reference to control CFU counts on agar without sorbic acid. At least three biological replicates were used in each experiment (exact numbers are clarified in figure legends). Each of the biological replicates (from independent experiments) were the average of three technical replicates for each sorbic acid concentration. Percentage survival data was then fitted to a lognormal distribution curve in R software as detailed below (Fitting to normal and lognormal distributions). Values derived with the curve fitting for IC_50_, heteroresistance and concentration at which 0.01% survival would be observed (MIC^*MODEL*^) were recorded.

### Inoculum-effect assay for heteroresistance

The inoculum effect (Steels et al. 2000) (Box 1) explains how experimentally measured MIC (MIC^*EXP*^) can increase with size of the cell inoculum, as increased cell number increases the chance that the inoculum includes phenotypically-variant cells that are hyper-resistant. Accordingly, the degree to which the inoculum effect alters MIC^*EXP*^ is related to the extent of heteroresistance: MIC^*EXP*^ would not be affected by inoculum size in a completely homogeneous cell population, whereas an impact of increasing inoculum size on MIC^*EXP*^ will be greater the broader the range of single-cell resistances (Figure 1A). It was reasoned that the magnitude of the inoculum effect could offer a novel, convenient measure of heteroresistance. Dose-response curves of growth-inhibitory effect typically follow a sigmoidal relationship (Stratford et al. 2013b) and have previously been modelled using Hill functions, (Geoghegan et al. 2020, Stratford et al., 2019), indicating a normal frequency distribution of single cell resistances. With that assumption, a relatively small number of MIC^*EXP*^ determinations at defined inoculum sizes becomes sufficient for curve-fitting (Figure 1B, top panel), to enable determination of standard deviation (σ) from the hypothetical normal distribution of cell resistances (Figure 1B, bottom panel). This standard deviation reflects the extent of heteroresistance, as if the difference between MIC^*EXP*^ values for different inoculum sizes increases, the normal distribution will have a broader bell curve, reflecting increased heteroresistance. In addition, this curve can be extrapolated to estimate the mean of the normal distribution, reflecting the population-median resistance (IC_50_, peak of the bell curve).

**Figure 1:**
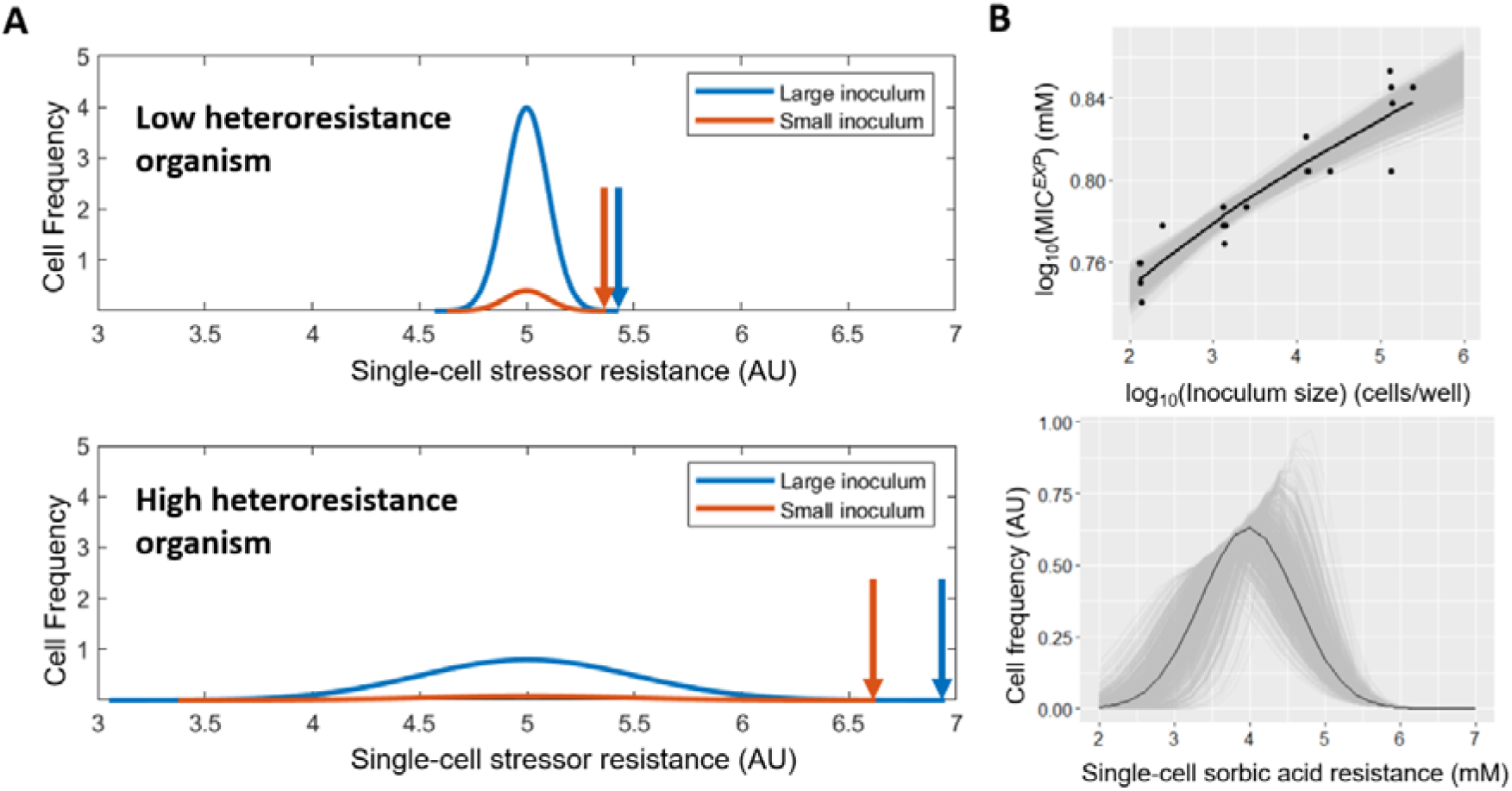
Establishing parameters for quantifying heteroresistance *via* the inoculum effect. **(A)** Modelled single-cell frequency distributions of stressor resistance (hypothetical) in low- and high-heteroresistance cell populations, at large and small inoculum sizes; these are modelled assuming single-cell resistances are normally distributed in the populations, with the total area under each curve equalling 1 and 0.1 for large and small inocula, respectively. Plots intersect the x-axis where survivor-cell frequency decreases below 0.001, corresponding to the theoretical point at which less than one cell of the relevant inoculum survives and equating to the MIC, indicated with an arrow. AU, arbitrary units. **(B)** Top panel: normal distribution curve fitted to MIC^*EXP*^ (Box 1) values for sorbic acid resistance (mM) at four optical densities (each with six replicates normalised to cells/well according to corresponding colony counts on agar) using inoculum effect methodology (see Methods) for strain 7812 (n=6 biological replicates). Black line, line of best fit. Grey lines, simulated normal distributions, with the distance between the upper and lower lines for each *x* value representing the 95% confidence interval for that point. Bottom panel: Simulation of frequency distribution of single cell resistances in strain 7812, based on normal distribution curve fitting in panel B (top).

Pre-culture of yeasts was carried out as described above. Exponential phase cells were diluted to OD_600_ ~0.2 (inoculum size in 75μl ~100,000 cells) and serially diluted three times by 10-fold, giving a lowest OD_600_ ~0.0002 (inoculum size in 75μl ~100 cells). Aliquots (75μl) of each inoculum size were transferred to wells of a flat-bottom, 96-well microplate (StarLab) and combined with 75μl of YEPD broth pre-supplemented with sorbic acid at double the desired, final sorbic acid concentration. After sealing with insulating tape, microplates were agitated at 600 rev/min for 60 s, placed in a sealed plastic bag, and incubated statically at 23°C for 21 days. The presence or absence of growth was noted visually under ~2X magnification using a magnifying glass. The lowest concentration at which no growth was detected in ≥1 of four technical replicates was scored as the experimental MIC (MIC^*EXP*^) for that inoculum from that biological replicate of the relevant isolate.

For inoculum effect measurements with the small panel of *Zygosaccharomyces* strains (Table S3), the procedure was carried out as above but using inoculum sizes of 10^5^, 10^4^, 10^3^ and 10^2^ cells/well, with concentration increments of 0.125 mM sorbic acid. For these particular assays, precise inoculum size was verified by plating 75μl aliquots of the OD_600_ ~0.0002 suspension (100 cells) on YEPD agar and incubating at 23°C for 7 days, the corresponding colony count was taken as the inoculum size of the lowest inoculum. The MIC^*EXP*^ results were fitted to a normal or lognormal distribution curve as described below.

### Fitting to normal and lognormal distributions

Royston et al. (1982) describe a formula to determine the expected value of the maximum of n random variables from a normal distribution (x_1_,..,x_n_), in terms of the theoretical mean (*μ*) and SD (*σ*) of that distribution (Eq.1).

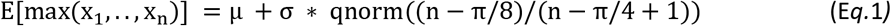

Assuming that the frequency distribution of antimicrobial resistance of n single cells from a homogeneous cell culture is approximated by a normal (or log-normal) one, this equation can be used to find the IC_50_ (the mean) and heteroresistance (the standard deviation) using data for MIC^*EXP*^ and their corresponding inoculum sizes (x).

Equation 1 was fitted to experimental data using non-linear least squares fitting, by using the nls() function in R, to return IC_50_ and heteroresistance values for each biological replicate. These were then used to calculate MIC^*MODEL*^ by taking the concentration value at the 99.99th percentile of the curve (where growth inhibition would occur in 99.99% of cells; Box 1). For each parameter, the mean and standard error of the mean (SEM) across biological replicates were taken.

Plotting of curves for MIC^*EXP*^ versus inoculum size was carried out using the ggplot package in R. Bootstraps to display 95% confidence interval of the line fitting were calculated using the Boot function in the ‘car’ package in R.

For dose-response data, sorbic acid concentration values and their corresponding percentage survival values were fitted to one minus a cumulative normal distribution function with mean and standard deviation equal to the IC_50_ and heterogeneity, respectively. Fitting was done using the nls() function in R software. Mean and SEM of each parameter were taken from the fitted curve as for dose response curves detailed above.

Normal or lognormal distribution curves fitted to data were displayed in R (ggplot package), in the case of inoculum effect data, and were displayed using the plot() function in MATLAB R20119b in the case of dose-response curve data. Box and whisker plots, bar charts and correlation plots were produced using Graphpad PRISM 9^TM^ software. Correlations containing points with error bars were produced using Microsoft Excel.

### Simulations

Simulation of hypothetical normal distributions for single cell resistances of organisms (for illustration of the inoculum effect concept) were produced using MATLAB R20119b. The Normal Probability Density Function (normpdf) was used to plot frequency distributions for low and high heteroresistance organisms (μ=5, σ=0.1 and μ=5, σ=0.1, respectively) with the frequency distribution divided by ten to simulate the small inoculum size.

### Data availability

All data are available in the present results section, in the supplemental material or on request. Genome sequence accession or sample numbers (ENA) are given above and in Table S1.

## Results

### Use of the inoculum effect for determining heteroresistance

In order to assess the contributions made by phenotypic heterogeneity (specifically, sorbic acid heteroresistance) and IC_50_ to the MIC, across large panels of species or strains, a new assay for measuring these parameters was developed based on the ‘inoculum effect’ (See methods; Steels et al. 2000), to allow higher throughput than traditional measurements of heteroresistance such as. from agar-based dose-response curve assays (Hewitt et al., 2016). Briefly, this method fitted a normal distribution curve of single cell resistances to experimental MIC (MIC^*EXP*^) values, by modelling MIC^*EXP*^ for a given inoculum as its corresponding point on the normal distribution curve (Figure 1B, top), e.g., MIC^*EXP*^ for 100 cells/well is the point were 1 in 100 cells would survive, MIC^*EXP*^ for 1000 cells/well is the point were 1 in 1000 cells would survive, etc. (see Methods). The median and standard deviation of the fitted normal distribution curve (Figure 1B, bottom) could then be taken as the IC_50_ and heteroresistance respectively.

For purposes of curve fitting, lognormal distributions were adopted as these gave better fit to dose response curve data than normal distributions (Figure S1). In addition, possible effect specifically of cell density (cell concentration) on the probability of growth occurring in a well was evaluated, e.g., due to stressor saturation or quorum sensing-type phenomena, as such effect could complicate interpretation of the inoculum effect which infers a total cell number effect. Only a minor possible effect of cell density was apparent, and similarly in high or low heteroresistance isolates (Figure S2). This result supported the inference that differences measured with inoculum effect methodology primarily reflected differences in heteroresistance rather than effects attributable to altered cell density.

To further assess the use of the inoculum effect method, IC_50_ (Figure 2A), heteroresistance (Figure 2B) and MIC (Figure 2C) determinations for six randomly selected strains from the *Zygosaccharomyces* panel (Table 1) were compared between the inoculum effect method and the dose-response curve method. The parameter MIC^*MODEL*^ was also taken, defined as the lowest concentration giving 99.99% inhibition of cell growth (Box 1) and consistent with previous publications in this field (Stratford et al. 2013b, 2014). This was calculated using the equation for the lognormal distribution curve fitted to MIC^*EXP*^ data (as in Figure 1B). For inoculum effect assays (carried out in broth), heteroresistance showed a moderately strong, significant correlation with heteroresistance values derived from dose-response curves (*r*=0.74), while IC_50_ and MIC^*MODEL*^ also showed strong correlations between the assays (*r*=0.853 and 0865, respectively). The inoculum effect assay was repeated on agar to assess possible effects of broth-versus agar-based assay formats (as in the above comparison) on the outcomes (Figure S3). Similar correlations were obtained as above indicating that for all the test parameters (IC_50_, heteroresistance and MIC^*MODEL*^), dose response results generally compare well with inoculum effect results obtained either in broth or on agar. These results supported use of the inoculum effect methodology, using a lognormal distribution for data, as an alternative heteroresistance assay to the established, but time-consuming dose-response curve method. Further assays below used the broth condition for inoculum effect assay, which offers higher throughput than on agar and is a condition also more relevant to the beverages and other product types in which sorbic acid is normally used.

**Figure 2:**
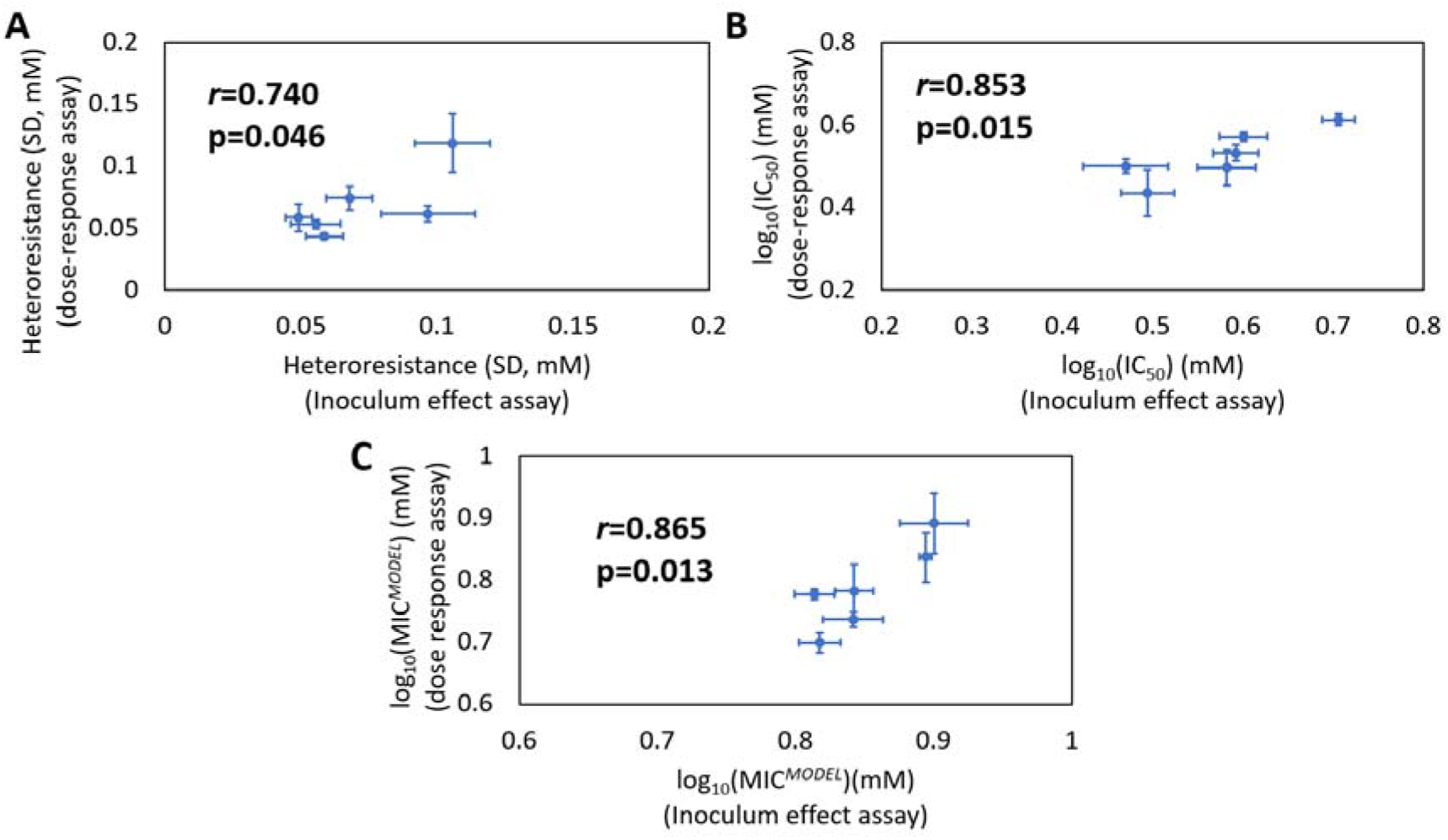
Corroboration of inoculum effect assay for measurement of IC_50_, heteroresistance and MIC^*MODEL*^. Comparison of parameter determinations for randomly selected *Z. parabailii, Z. bailii* and *Z. pseudobailii* isolates (isolates 7788, 7812, 7851, 7829, 7769, 7842). Values for IC_50_ **(A)**, heteroresistance **(B)** and MIC^*MODEL*^ **(C)** were derived either from dose-response curve on agar or inoculum effect assay in broth (using MIC^*EXP*^ values at two inoculum sizes, see Methods). Points represent means from 6 biological replicates for each isolate +/- SEM, with *r* values calculated using linear regression and p values were calculated using one-tailed linear regression.

### High-throughput determination of survival parameters for *Zygosaccharomyces* isolates

To investigate relationships between heteroresistance or IC_50_ with MIC^*MODEL*^, we applied the inoculum effect methodology to a panel of *Zygosaccharomyces* sp. Isolates. To select strains for study, an initial panel of 111 yeasts isolated from foods (see Table S1) were whole-genome sequenced (WGS); most of these had previously been considered as *Z. bailii* isolates (Stratford et al. 2013b, 2014, 2019). The sequencing revealed that several isolates previously characterised as *Z. bailii* were in fact *Z. parabailii* or *Z. pseudobailii* (see Methods). These species are known interspecies hybrids formed between *Z. bailii* and unknown, closely related ancestors. From these 111 isolates, for further study a panel of 29 strains were selected, to include isolates sampled from across the span of the phylogenetic tree (Figure S5) and diverse sources of isolation. This ‘Main *Zygosaccharomyces* panel’ comprised 5, 17 and 7 isolates of *Z. bailii, Z. parabailii* and *Z. pseudobailii* respectively (Table 1). For parameter measurement (heteroresistance, IC_50_ and MIC^*MODEL*^) inoculum effect methodology was used, with curves fitted to MIC^*EXP*^ values taken at 10^5^ and 10^2^ cells/well. There was considerable variation in heteroresistance, IC_50_, and MIC^*MODEL*^ among isolates of the panel, with ranges of approximately 0.03–0.11, 2.3–5.4 and 4.3–9.2 mM for heteroresistance, IC_50_ and MIC^*MODEL*^, respectively (Table 1; Figure 3A). Analysis for each subspecies indicated that *Z. pseudobailii* had a significantly greater mean heteroresistance than *Z. parabailii* and *Z. bailii*, whereas the mean IC_50_ for *Z. pseudobailii* was significantly lower than for *Z. parabailii* (Figure 3B). There were no significant differences in average MIC^*MODEL*^ between the different subspecies (two sample t-test assuming unequal variances), although there was both a broader spread of values and relatively low mean for *Z. bailii* isolates (Figure 3B).

**Figure 3:**
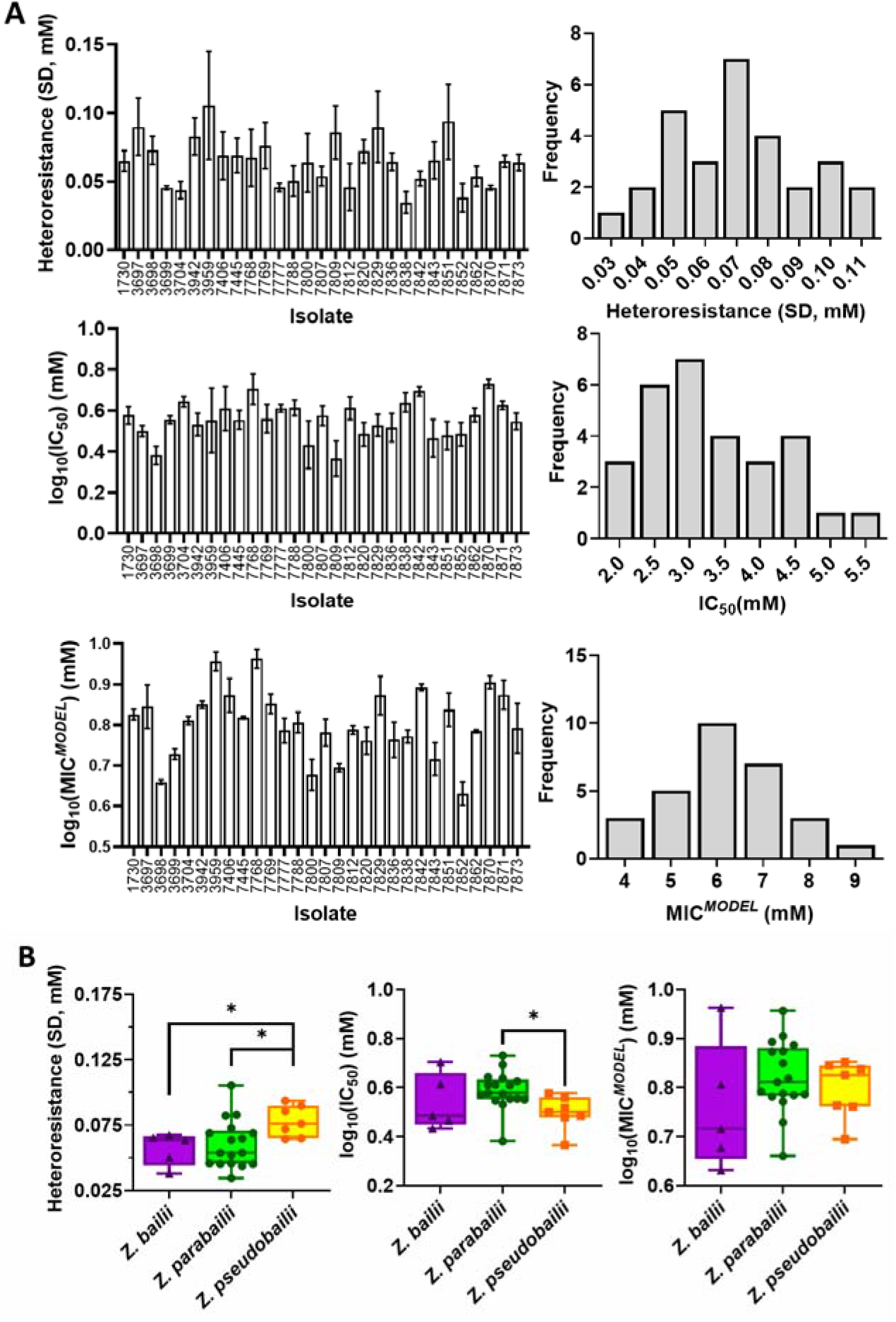
Heteroresistance, IC_50_ and MIC^MODEL^ across a panel of 29 *Zygosaccharomyces* sp. isolates grown with sorbic acid. **(A)** Left panels: heteroresistance, IC_50_ and MIC^*MODEL*^ determinations for individual strains. Values are means ± SEM of average of parameters for three curves independently fitted to each biological replicate (MIC^*EXP*^ values measured at two inoculum sizes.) Right panels: Frequency histograms showing spread of heteroresistance, IC_50_ and MIC^*MODEL*^ in the 29-strain panel (strain list in Table 1). Values on the x axis indicate centre of each histogram bin, where divisions between bins were at the midpoints between adjacent values shown. **(B)** Data adapted from (A) to compare the distributions of heteroresistance, IC_50_ and MIC^*MODEL*^ among the isolates of the three subspecies. Significant differences are indicated (*p<0.05), using two-sample *t*-test assuming unequal variances (n =5, 17 and 7, for *Z. bailii, Z. parabailii* and Z. *pseudobailii*, respectively).

To further investigate the relationships between heteroresistance, IC_50_ and MIC^*MODEL*^, the correlation between these parameters was calculated. Across the main *Zygosaccharomyces* panel of 29 strains, strains with high IC_50_ tended to have high MIC^*MODEL*^ and *vice versa* (*r*=0.675, p<0.0001), whereas heteroresistance was not significantly correlated with MIC^*MODEL*^ (*r*=0.326, p=0.08) (Figure 4A). Unexpectedly, strains with a high IC_50_ tended to have a low heteroresistance (*r*=0.586, p<0.01) (Figure 4A). When the subspecies were examined independently, *Z. bailii* (Figure 4B), *Z. parabailii* (Figure 4C) and *Z. pseudobailii* Figure 4D) also each showed a significant correlation between IC_50_ and MIC^*MODEL*^ and no significant correlation between heteroresistance and MIC^*MODEL*^. In the case of *Z. parabailii*, it was noted that calculated correlations were strongly influenced by a single isolate (3698) with a much lower IC_50_ than all the other *Z. parabailii* isolates (Figure 4C). When the *Z. parabailii* dataset was analysed without this single isolate (n=16), heteroresistance was strongly correlated with MIC^*MODEL*^ (*r*= 0.660, p<0.01) and IC_50_ was not (*r*= 0.265, p=0.32) (Figure S4). The correlation between heteroresistance and MIC^*MODEL*^ in this sample could suggest that heteroresistance is more strongly related to MIC^*MODEL*^ when variation in IC_50_ is relatively limited, as was the case in *Z. parabailii* after removal of isolate 3698 (see Discussion).

**Figure 4:**
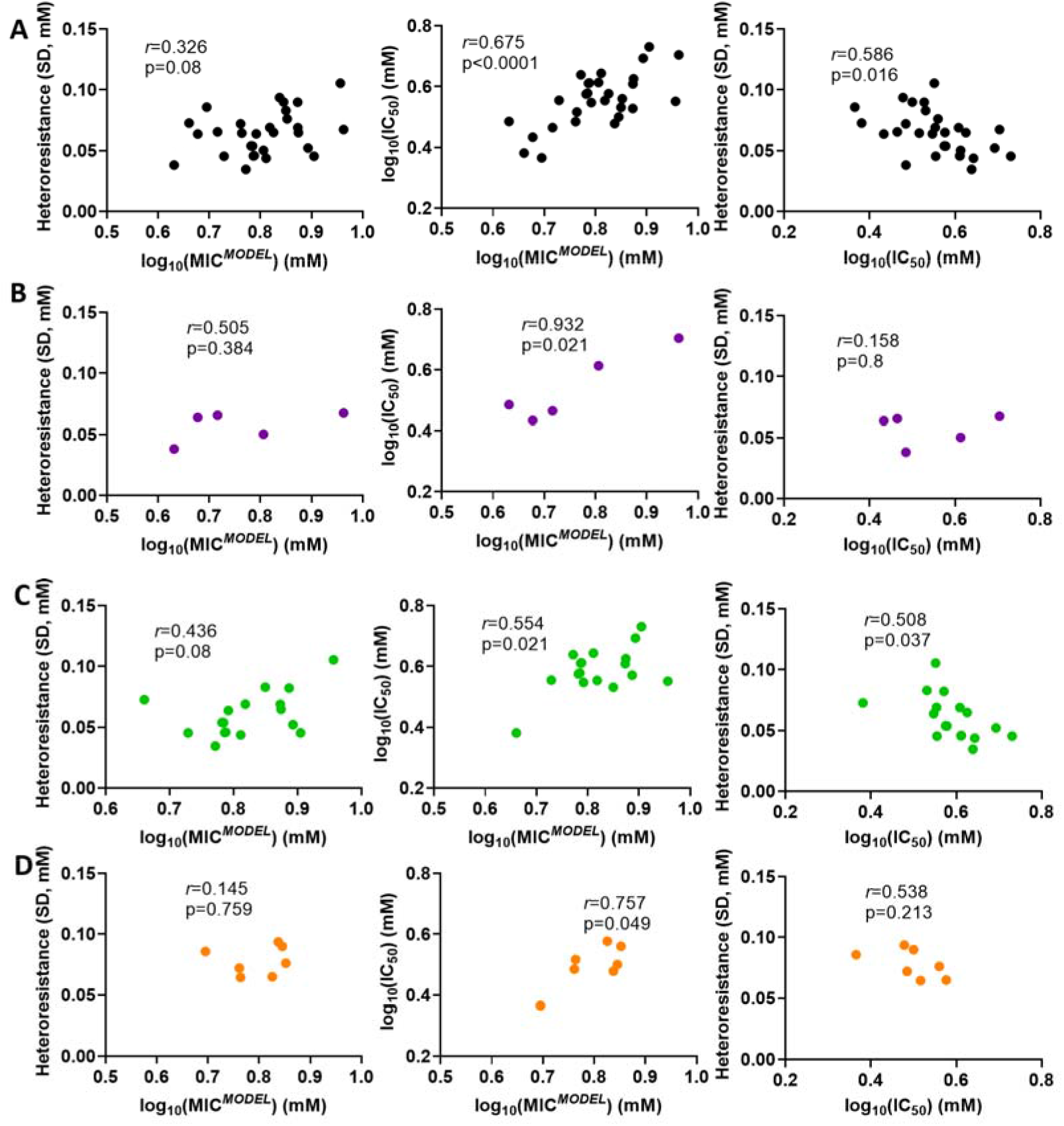
Relationships between heteroresistance, IC_50_ and MIC^*MODEL*^ in the main panel of *Zygosaccharomyces* sp. isolates. Data were obtained as described in Figure 3. **(A)** Cross-species panel of *Zygosaccharomyces* isolates (n=29) (Table 1). **(B)** *Z. bailii* only (n=5). **(C)** *Z. parabailii* only (n=17). **(D)** *Z. pseudobailii* only (n=7).

### Trends in heteroresistance, IC_50_ and MIC^*MODEL*^ across a wide panel of diverse yeast species

To investigate if the trends in sorbic acid heteroresistance described above may be replicated in other spoilage yeast species, dose response curve data from a ‘wide panel’ of spoilage yeasts isolates (Table S2) produced in a previous survey were analysed. These isolates encompassed 26 species from 12 different yeast genera. Three *Z. parabailii* and two *S. cerevisiae* strains were present in the panel, whereas all other species were represented by one isolate only. To avoid mixing intra- with inter-species comparisons, values averaged across the three *Z. parabailii* and across the two *S. cerevisiae* isolates were used for these two species. This wide yeast panel produced a notably wider range of IC_50_ and MIC^*MODEL*^ values than the main *Zygosaccharomyces* panel (Figure 5A,B). This was primarily due to comparatively low IC_50_ and MIC^*MODEL*^ values for certain species of the wide panel, such as *Rhodotorula mucilaginosa* (IC_50_~0.337mM, MIC^*MODEL*^~0.467 mM) and *Zygosaccharomyces rouxii* (IC_50_~1.34mM, MIC^*MODEL*^~2.45mM), two species previously associated with food environments where sorbic acid is not used (Pitt & Hocking, 2009). The two panels exhibited similar variation in heteroresistance between isolates, but the *Zygosaccharomyces* isolates showed significantly higher mean heteroresistance (p<0.0001) (Figure 5B). IC_50_ was very strongly correlated with MIC^*MODEL*^ in the wide panel (*r*=0.986, p<0.0001) (Figure 5C). The wide panel also showed a weaker correlation between heteroresistance and MIC^*MODEL*^ (*r* = 0.487, p < 0.05). There was a weak, non-significant correlation between heteroresistance and IC_50_, which was weaker than that with MIC^*MODEL*^ suggesting that the relationship between heteroresistance and MIC^*MODEL*^ was not simply a reflection of the strong relationship between IC_50_ and MIC^*MODEL*^. The results collectively indicate a close relationship between MIC and population-median resistance (IC_50_) and additionally, albeit relatively weakly, with heteroresistance, as discussed below.

**Figure 5:**
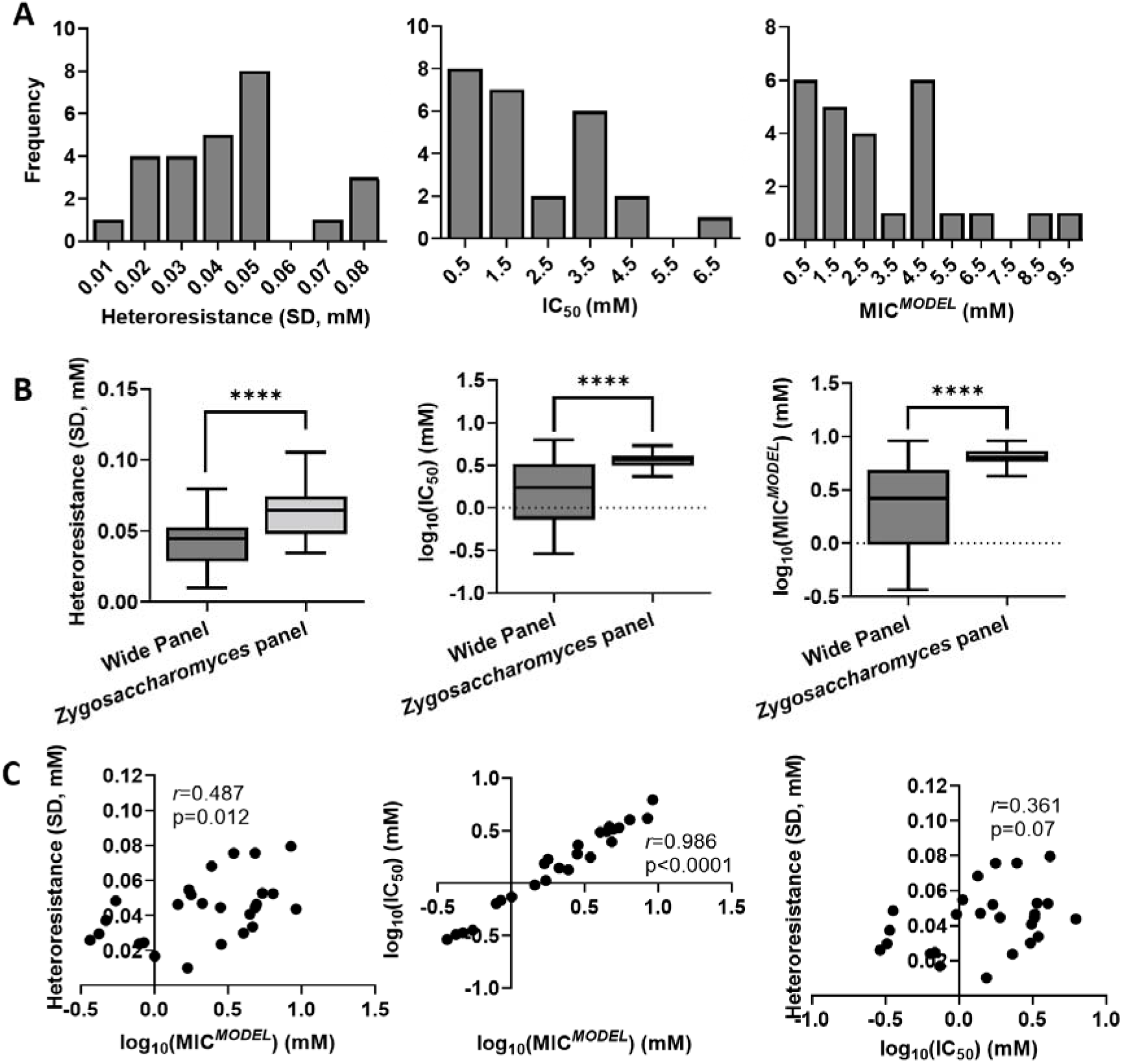
Trends in sorbic acid heteroresistance, IC_50_ and MIC^*MODEL*^ in a wide panel of diverse yeast species. **(A)** Frequency histograms showing the spread of heteroresistance, IC_50_ and MIC^*MODEL*^ measured using the dose response curve method. Values for individual 26 species are listed in Table S2. Values for individual isolates were means from three biological replicates. **(B)** Box and whisker plots comparing the parameters between the wide panel (n=26) and the *Zygosaccharomyces* panel (n=29) data from Figure 4. ***, p<0.0001 according to a two-sample *t*-test assuming unequal variances. **(C)** Correlations between heteroresistance, IC_50_ and MIC^*MODEL*^ for the wide panel.

## Discussion

This study showcases the relevance of IC_50_ and heteroresistance parameters for an inhibitor’s MIC against organisms, here focussing on the preservative sorbic acid. The novel, high-throughput inoculum effect methodology developed here allowed heteroresistance to be determined in a large number of microorganisms, in a manner less laborious than an established method like dose-response assay on agar. In addition, the microplate format of the new assay demands only modest physical incubator space and is amenable to automation. This methodology offers the potential for screens of heteroresistance in new ecological or industrial contexts, for example in modelling and quantitative risk assessments in the food industry. Our results indicate that a dominant indicator of differences in MIC between strains is differences in IC_50_, with heteroresistance (e.g. incidence of rare, hyper-resistant cells) playing a smaller role.

The experimental data were consistent with lognormal distributions of single-cell resistances in the yeast cell populations. Hill functions have previously been employed to model dose response curves using log10 values for stressor concentration (Geoghegan et al., 2020; Stratford et al. 2019), a model which assumes single-cell resistances are lognormally distributed. Here, while the fit of the lognormal distribution curve to the data was only marginally better than a normal distribution, for this study we were more interested in differences in relative distributions of single cell resistances, therefore lognormal was suited to the study’s aims.

The possibility that cell-concentration may make a contribution to the inoculum effect that is independent of effects due to resistant-cell incidence was investigated, e.g., arising from stressor saturation or quorum sensing-type phenomena, for example. Some previous reports of the inoculum effect have not made this distinction (Loffredo et al. 2020, Smith et al., 2018). However, Scheler et al. (2020) demonstrated that inoculum effect in *E. coli* for cefotaxime resulted from heteroresistance rather than cell concentration, while Steels et al. (2000) ruled out certain effects of cell concentration (e.g. altered adsorption) in the inoculum effect of *Z. bailii* with sorbic acid. On the other hand, cell concentration affected survival probability of heat-shocked *S. cerevisiae*, attributed to cooperative glutathione excretion at high cell density (Laman Trip & Youk, 2020). To our knowledge, no such mechanisms have been reported for sorbic acid stress. Here, analyses indicated that cell concentration may make only a minor contribution to the increasing sorbic acid MIC^*MODEL*^ as inoculum size was increased (between 10^2^ and 10^4^ cells per well). One possible explanation for that small effect could be related to the documented metabolism of sorbic acid by *Z. bailii* (Mollapour and Piper, 2001).

Heteroresistance, IC_50_ and MIC^*MODEL*^ values obtained on agar (from dose response assay) showed significant correlation with the same parameters obtained in broth (from inoculum effect assay). This consistency for all three parameters whether measured in broth or agar is in line with work elsewhere reporting similar resistances to other stressors in broth versus agar (Morris et al., 2020; Albano et al., 2021; Fuchs et al., 2021). Robustness of the heteroresistance phenotype to the form of growth environment is consistent with heteroresistance being an intrinsic property of cell populations, not strongly affected by environment. This property allows resistant subpopulations to arise independent of stressor exposure, for example, so enabling pre-priming for possible future perturbations. Such pre-priming of heteroresistance has been described for other stressor scenarios (Jing et al. 2018, Levy et al., 2012; Stratford et al. 2014), and is in line with understanding of the wider premise for phenotypic heterogeneity, or bet-hedging (Avery, 2006; Ackermann, 2015).

There was considerable variation in all three parameters between strains and species of the *Zygosaccharomyces* genus and across a wider panel of spoilage yeast species. *Z. bailii* is considered as more heteroresistant to sorbic acid than *S. cerevisiae* (Stratford et al. 2014). Across the main *Zygosaccharomyces* panel and the constituent subspecies, IC_50_ was more clearly correlated with MIC^*MODEL*^ than was heteroresistance. However, in the case of *Z. parabailii* this trend was reversed by the removal of a single outlying strain with an uncharacteristically low IC_50_ value. The removal of this outlier considerably reduced the variation in IC_50_ for the subspecies (standard deviation reduced to 0.055 from 0.076) but had negligible impact on variation in heteroresistance (standard deviations 0.0188 and 0.0186). This could suggest that relationships between heteroresistance and MIC are masked when IC_50_ variation is high and potentially dominating effects on MIC across different organisms. Accordingly, heteroresistance may be more important for MIC in species or contexts that support lower variation in IC_50_.

Regarding the question of why one (sub-)species might evolve a higher average preservative heteroresistance than others (e.g. *Z. pseudobailii*, Figure 3B), drivers could be related to the organisms’ ecologies or relative incidences. This may include the size of inoculum typically associated with spoilage incidence of an organism, as larger inocula could amplify the effect that heteroresistance has on the MIC. Alternatively, the frequency of transition of the growth environment from preservative-rich to -poor, as frequent changes are thought to favour high heterogeneity (Avery, 2006). Modelling of gene expression patterns has indicated that gene expression noise confers a stronger fitness advantage when population-averaged gene expression is further from the optimal level (Duveau et al. 2018), a situation more often encountered in fluctuating environments. Regarding the particular example of this study, a less stable niche for *Z. pseudobailii* is partially supported by this organism being found primarily in viscous foodstuffs (Table 1), although a similar argument could be made for *Z. parabailii* (where average heteroresistance was lower) which colonises solid as well as liquid foodstuffs.

There was a negative correlation between heteroresistance and IC_50_ across the main *Zygosaccharomyces* panel. One suggestion to explain this might be that heteroresistance is more strongly selected in low-IC_50_ strains in compensation for a disparity between their population-median resistance (IC_50_) and optimal resistance when faced with stressor. That would be in line with other previously described scenarios, including the example mentioned above where noisy gene expression is especially beneficial when average gene expression is far from the optimum (Duveau et al. 2018).

Other trends were evident here across a wider panel of spoilage yeast species. In particular, a weak but significant positive correlation between heteroresistance and MIC emerged. This suggests that heteroresistance may make a larger contribution to MIC differences between distantly related species of spoilage yeasts than across individual isolates of the *Zygosaccharomyces* genus. At the same time, a correlation between IC_50_ and MIC in the wide panel was stronger than in the main *Zygosaccharomyces* panel, possibly reflecting the wider range of IC_50_ phenotypes across the wide panel. While specific trends were different between the two panels the overall conclusion was similar, that IC_50_ is more closely related than heteroresistance to preservative MIC.

Why heteroresistance has not been strongly associated with sorbic acid resistance in this study requires consideration. One explanation is that if the metabolic cost to a cell of resisting sorbic acid stress is low, the fitness benefit of bet-hedging over increasing population-average resistance would be reduced. While some strategies for sorbic acid resistance such as increased expression of efflux pumps (Geoghegan et al. 2020) would be metabolically costly, favouring heteroresistance (Jing et al. 2018), other proposed resistance mechanisms such as potential maintenance of a more fermentative metabolism (Stratford et al. 2020) might confer resistance at a comparably low cost, mitigating the fitness benefit of heteroresistance over IC_50_. It is possible that stronger heteroresistance phenotypes would be found in niches where the metabolic cost of pre-priming (all) cells for stress was greater. A possibly-related observation was overrepresentation of the high-IC_50_, low-heteroresistance species *Z. parabailii* in the initial *Zygosaccharomyces* panel used in this study (n=71, 19 and 21 for *Z. parabailii*, *Z. pseudobailii* and *Z. bailii* respectively, Table S1). As this reflects isolation of *Z. parabailii* from a larger number of food spoilage events, it is consistent with the suggestion that a high IC_50_ as opposed to high heteroresistance may be a more effective evolutionary strategy for growing in preserved foods, though of course we cannot discount that other traits of *Z. parabailii* account for its preponderance. It should also be noted that this study was limited to food spoilage isolates only and other relationships may be manifest in other ecological niches.

### Conclusion

We capitalised on the inoculum effect principle to develop a new, high-throughput method for assaying heteroresistance. Application of this assay to diverse spoilage yeasts revealed that heteroresistance and especially IC_50_ are related to MIC, whereas a relationship with heteroresistance was not observed when considering a panel only of *Zygosaccharomyces* sp. isolates. Overall, IC_50_ appears to be a strong indicator of MIC, but heteroresistance may also be related to MIC between distantly related species or in contexts where IC_50_ variation is low. This work highlights certain limitations of conventional MIC measurements as, due to the effect of heteroresistance, the apparent resistance of an organism could vary markedly according to inoculum size. The average resistance of a cell population, its heteroresistance and the inoculum size should all be considered in selecting appropriate antimicrobial concentrations for specific applications, e.g., food preservation. In experimental work (e.g. in microbial challenge testing) this would translate to using an adequate population size (e.g. 10^4^ cells) in the desired test volume.

## Acknowledgements

This work was supported by the Biotechnology and Biological Sciences Research Council (grant number B/T508949/1). We thank Malcolm Stratford for kindly sharing yeast isolates and data.

